# Postsynaptic neuroligin-1 mediates presynaptic endocytosis during neuronal activity

**DOI:** 10.1101/2021.07.20.453031

**Authors:** Jiaqi Keith Luo, Holly Melland, Jess Nithianantharajah, Sarah L Gordon

## Abstract

Fast, high-fidelity neurotransmission and synaptic efficacy requires tightly regulated coordination of pre- and postsynaptic compartments and alignment of presynaptic release sites with postsynaptic receptor nanodomains. Neuroligin-1 (Nlgn-1) is a postsynaptic cell-adhesion protein exclusively localised to excitatory synapses that is crucial for coordinating the transsynaptic alignment of presynaptic release sites with postsynaptic AMPA receptors as well as postsynaptic transmission and plasticity. However, little is understood about whether the postsynaptic machinery can mediate the molecular architecture and activity of the presynaptic nerve terminal, and thus it remains unclear whether there are presynaptic contributions to Nlgn1-dependent control of signalling and plasticity. Here, we employed a presynaptic reporter of neurotransmitter release and synaptic vesicle dynamics, synaptophysin-pHluorin (sypHy), to directly assess the presynaptic impact of loss of Nlgn1. We show that lack of Nlgn1 had no effect on the size of the readily releasable or entire recycling pool of synaptic vesicles, nor did it impact exocytosis. However, we observed significant changes in the retrieval of synaptic vesicles by compensatory endocytosis, specifically during activity. Our data extends growing evidence that synaptic adhesion molecules critical for forming transsynaptic scaffolds are also important for regulating activity-induced endocytosis at the presynapse.

## Introduction

Activity-dependent neural signalling and plasticity relies on the coordination of pre- and postsynaptic compartments. The alignment of presynaptic release sites with postsynaptic receptor nanodomains is important for tightly regulating synaptic efficacy and for fast, high fidelity neurotransmission. This critically relies on transsynaptic organization by adhesion molecules such as the neurexin-neuroligin complex (Sudhof, 2008;Haas et al., 2018). Pre-synaptic neurexin drives active zone formation (Missler et al., 2003) and postsynaptic neuroligin-1 (Nlgn-1) recruits scaffolds, such as PSD-95, and NMDA and AMPA receptors (Graf et al., 2004;Heine et al., 2008;Mondin et al., 2011;Budreck et al., 2013).

Nlgn-1 is the only member of the neuroligin family of synaptic cell-adhesion proteins (Nlgn1-4) that is exclusively localised to the postsynaptic membrane of excitatory synapses (Song et al., 1999). Nlgn1 is a single pass transmembrane protein with an extracellular catalytically inactive acetylcholine esterase domain and an intracellular c-terminus which interacts with scaffolding proteins such as PSD-95 (Ichtchenko et al., 1995;Irie et al., 1997). It forms a constitutive homo- or heterodimer (with its family member neuroligin-3, Nlgn3), and via its extracellular N-terminal domain binds to two presynaptic neurexin molecules in a calcium-dependent manner to mediate trans-synaptic cell adhesion (Nguyen and Sudhof, 1997;Arac et al., 2007). Whilst Nlgn1, and other Nlgn family members, are not required for formation of the synaptic ultrastructure (Varoqueaux et al., 2006), Nlgn1 is important for coordinating the alignment of presynaptic release sites with postsynaptic AMPA receptors that is required for synaptic efficacy (Haas et al., 2018). Importantly, Nlgn1 has been consistently shown to be crucial for postsynaptic transmission and plasticity mediated by NMDA receptors (NMDAR). NMDAR-mediated EPSC amplitude and NMDA/AMPA ratio are robustly reduced in both *in vitro* and *in vivo* Nlgn1 knockdown or knockout systems (Chubykin et al., 2007;Kim et al., 2008;Blundell et al., 2010;Jung et al., 2010;Budreck et al., 2013;Espinosa et al., 2015;Jiang et al., 2017)

Whether there are presynaptic drivers that contribute to the signalling and plasticity changes induced by loss of Nlgn1 remains unclear, as relatively little has been done to examine whether the postsynaptic machinery can mediate the molecular architecture and activity of the presynaptic nerve terminal. It has recently been shown that overexpression of Nlgn1 increases the number of synaptic vesicles undergoing exocytosis in response to a 20Hz stimulus train (van Stegen et al., 2017). Additionally, loss of Nlgn1 has no effect on release probability or rate of exocytosis, but does decrease the degree to which presynaptic nerve terminals take up a fluorescent marker of cycling vesicles, suggesting perturbation of the synaptic vesicle cycle (Wittenmayer et al., 2009). Absence of α- and β-neurexins also reduces the size of the readily releasable pool of vesicles, as does expression of truncated neurexin which is unable to bind its postsynaptic partners (Quinn et al., 2017). Collectively, these data suggest that the transsynaptic binding of Nlgn1 with neurexin may be important for modulating presynaptic activity and neurotransmitter release. However, whether Nlgn1 is necessary for specific processes in the synaptic vesicle cycle remains an open question.

Here, we employed a presynaptic reporter of neurotransmitter release and synaptic vesicle dynamics, synaptophysin-pHluorin (sypHy), that allowed us to directly assess the impact of loss of Nlgn1 on the functioning of the nerve terminal in a manner that does not rely on measures of postsynaptic signalling. Surprisingly, we found that absence of Nlgn1 leads to changes in the retrieval of synaptic vesicles by compensatory endocytosis, specifically during activity.

## Materials and Methods

### Materials

Synaptophysin-pHluorin (sypHy) was provided by Prof Leon Lagnado (University of Sussex, UK). Neurobasal media, B-27 supplement, L-glutamine, penicillin/streptomycin, Minimal Essential Medium (MEM), Dulbecco’s Modified Eagle Medium (DMEM), Lipofectamine 2000 were obtained from Thermo Fisher Scientific. Bafilomycin A1, 6-cyano-7-nitroquinoxaline-2,3-dione and DL-2-Amino-5-phosphonopentanoic acid were obtained from Sapphire Bioscience. All other reagents were obtained from Sigma-Aldrich.

### Hippocampal neuronal cultures

All procedures were approved by the Florey Animal Ethics Committee and performed in accordance with the guidelines of the National Health and Medical Research Council Code of Practice for the Care and Use of Animals for Experimental Purposes in Australia. Heterozygous *Nlgn*^+/-^ mice were obtained from Prof. Nils Brose (Varoqueaux et al., 2006) and backcrossed on a C57BL/6 background as described (Luo et al., 2020). *Nlgn1*^-/-^ mice and WT littermate controls were generated at The Florey by mating *Nlgn*^+/-^ females and males. Mouse colonies were maintained in a temperature controlled (≈ 21°C) room on a 12 h light/dark cycle (lights on at 07:00) and group housed in individually ventilated cages with food and water available *ad libitum*. WT x WT and *Nlgn1*^-/-^ x *Nlgn1*^-/-^ mice were time mated overnight and visualisation of a vaginal plug on the following morning was considered as embryonic day (E) 0.5. Dissociated primary hippocampal enriched neuronal cultures were prepared from E16.5-18.5 WT and *Nlgn1*^-/-^ mouse embryos of both sexes by trituration of isolated hippocampi to obtain a single cell suspension, which was plated at a density of 5×10^4^ cells/coverslip on poly-D-lysine and laminin-coated 25 mm coverslips. Cultures were maintained in neurobasal media supplemented with B-27, 0.5 mM L-glutamine and 1% v/v penicillin/streptomycin. After 72 hours cultures were further supplemented with 1 μM cytosine β-d-arabinofuranoside to inhibit glial proliferation. Cells were transfected after 7 days in culture with Lipofectamine 2000 as described (Gordon et al., 2011). In all experiments cells were co-transfected with 1µg each of mCherry (as a transfection marker) and sypHy. Cells were imaged after 13-15 days in culture.

### Fluorescence imaging

Hippocampal cultures adhered to coverslips were mounted in a Warner imaging chamber with embedded parallel platinum wires (RC-21BRFS) and placed on the stage of Leica SP8 confocal microscope for determining surface expression of sypHy and synaptic vesicle pool size, and Zeiss Axio Observer 7 epifluorescence microscope for experiments assessing exocytic and endocytic dynamics. Neurons expressing sypHy were visualized with a x40 oil immersion objective (NA 1.3) at 475nm (Zeiss) or 488nm (Leica) excitation wavelength with a GFP filter. Z-stacks were acquired on the confocal fluorescence microscope (Leica) which included the entire volume of presynaptic terminals. Nyquist sampling was used for calculation of z-step size. Neurons were subjected to continuous perfusion with imaging buffer (in mM: 136 NaCl, 2.5 KCl, 2 CaCl2, 1.3 MgCl2, 10 glucose, 10 HEPES, pH 7.4) throughout image acquisition unless noted otherwise. Electrical stimulations (100 mA, 1ms pulse width; 60 action potentials delivered at 30 Hz, 1200 action potentials delivered at 10 Hz, or 300 action potentials at 10 Hz) were delivered in the presence of 10 μM 6-cyano-7-nitroquinoxaline-2,3-dione and 50 μM DL-2-Amino-5-phosphonopentanoic to inhibit network activity. 1 μM bafilomycin A1 was added to the imaging buffer to inhibit reacidification of synaptic vesicles and allow assessment of synaptic vesicle pools and exocytosis. At the final stage of image acquisition, neurons were perfused with alkaline imaging buffer (50 mM NH4Cl substituting 50 mM NaCl) to reveal total sypHy fluorescence. All experiments were conducted at 37°C.

### Measuring vesicular sypHy

The cell surface fraction of sypHy was revealed by bathing neurons in imaging buffer, and then perfusing with acidic imaging buffer (20 mM 2-(N-morpholino) ethanesulfonic acid substituted for HEPES, pH 5.5) to quench cell surface sypHy fluorescence and retain background autofluorescence. Neurons were washed in imaging buffer, and then exposed to alkaline imaging buffer to reveal total sypHy fluorescence. The vesicular fraction of sypHy as a percentage of total was calculated as ([(total – background) – (surface – background)] / (total – background)) x100). The second period of normal imaging buffer perfusion was taken as the baseline.

### Data analysis

Regions of interest of identical size (within each experiment) were placed over presynaptic terminals using FIJI to measure fluorescence intensity. Terminals which did not respond to stimulation or whose maximum fluorescence falls outside the period of alkaline buffer perfusion were excluded. Details of quantification of fluorescence imaging can be found in supplemental methods. Curve fitting was performed in MATLAB R2019. Statistical testing was performed using GraphPad Prism 8. Unpaired t-tests were used for hypothesis testing where response did not deviate significantly from normality and Welch’s t-test was used where variances differed significantly between genotypes. N indicates individual field of view from an independent coverslip, sampled from at least 3 independent cultures for each experiment.

## Results

To investigate whether Nlgn1 can act in a transsynaptic manner to impact presynaptic function, we employed synaptophysin-pHluorin (sypHy), a genetically encoded pH-sensitive fluorescent protein which has been extensively used to characterise synaptic vesicle dynamics. The pHluorin moiety is tagged to the lumenal domain of a synaptic vesicle protein (synaptophysin); its fluorescence is quenched at rest by the acidic intravesicular pH and increases upon exocytosis as the vesicle fuses with the plasma membrane and the pHluorin is exposed to the neutral extracellular environment (Fig 1A). sypHy fluorescence decreases as the vesicle membrane and protein cargo are retrieved by compensatory endocytosis and vesicles reacidify, but this can be blocked by the use of the vATPase inhibitor bafilomycin (Fig 1A). Perfusion with NH4Cl reveals the total sypHy fluorescence in the presynaptic terminal. To probe the contribution of Nlgn1 to synaptic vesicle dynamics, sypHy was transfected into cultured hippocampal neurons prepared from embryonic wildtype (WT) or null mutant mice lacking Nlgn1 (*Nlgn1*^-/-^).

**Figure 1.**
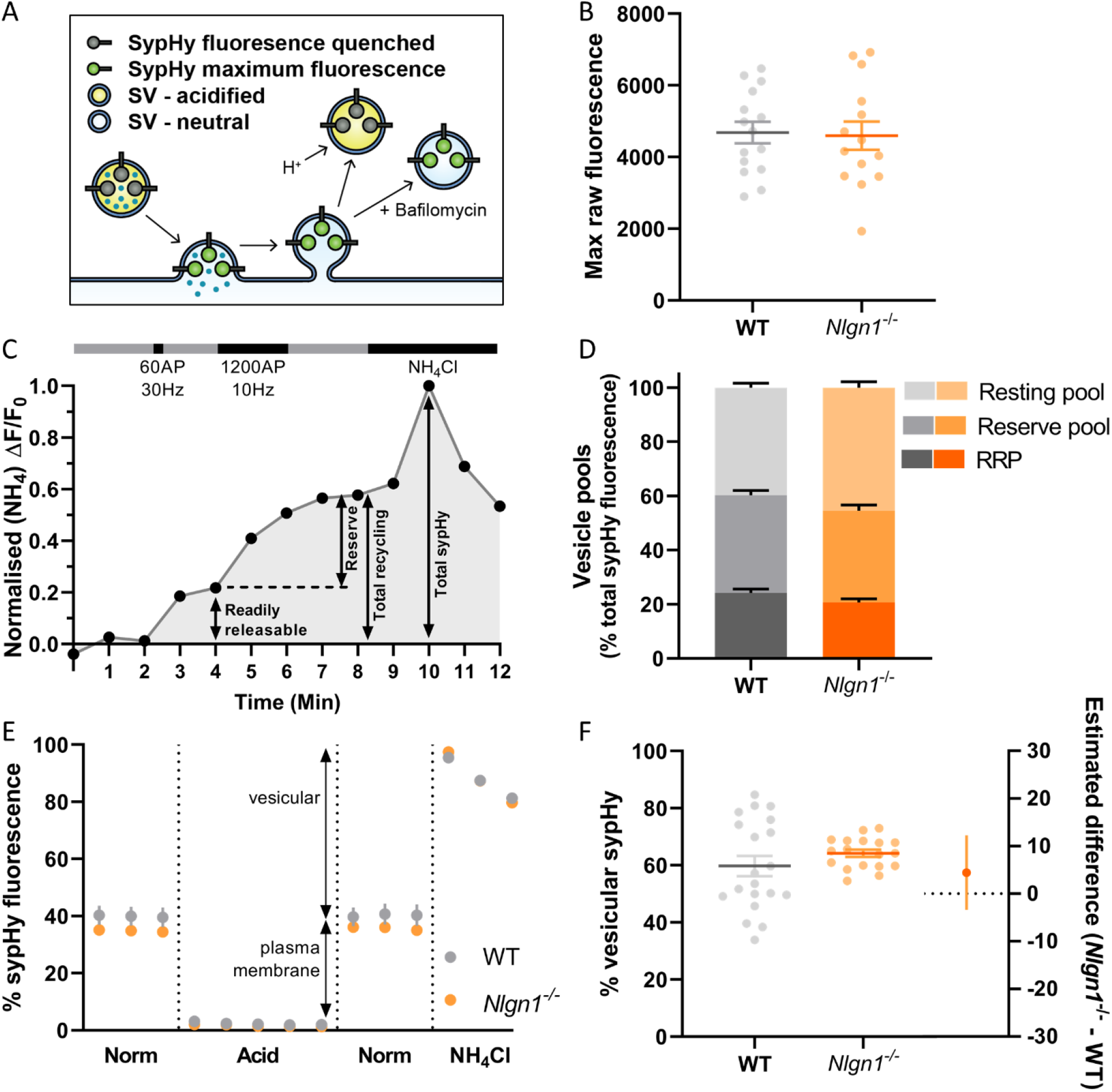
(over page): Lack of Nlgn1 has does not impact the size of synaptic vesicle pools. *Nlgn1*^-/-^ and WT cultured neurons were transfected with sypHy. **A**: sypHy is a genetically encoded, pH-sensitive fluorescence reporter of synaptic vesicle (SV) dynamics. **B**: Scatter plot of total sypHy fluorescence in *Nlgn1*^-/-^ and WT neurons, as assessed by perfusion with NH4Cl; mean ± SEM indicated. N = 14-15 individual fields of view from independent coverslips, unpaired t-test, p = 0.8601. **C**: Stimulation protocol for assessing specific synaptic vesicle pools. Neurons were stimulated with 60 action potentials (AP) at 30Hz to trigger fusion of the readily releasable pool (RRP), followed by 1200 AP at 10Hz to mobilise the reserve pool of vesicles (together composing the total recycling pool), in the presence of bafilomycin to inhibit vesicle reacidification. The remaining vesicles, which do not undergo fusion, comprise the resting pool. Fluorescence change was monitored over time, and neurons were then perfused with NH4Cl to reveal total sypHy fluorescence; fluorescence change from baseline was normalized to the total sypHy fluorescence. **D**: Synaptic vesicle pool sizes in *Nlgn1*^-/-^ and WT synapses. Size of distinct vesicle pools (readily releasable, reserve and resting as a proportion of total sypHy fluorescence) averaged over individual coverslips are shown in stacked column graphs. N = 10 – 13 individual fields of view from independent coverslips, values represent mean ± SEM. Two-way ANOVA with Sidak’s multiple comparison test; RRP p = 0.4256, reserve pool p = 0.7539; resting pool p = 0.0744. **E**: Fluorescence trace for quantifying the partitioning of sypHy to distinct membranous compartments in *Nlgn1*^-/-^ and WT synapses. Neurons transfected with sypHy were perfused with normal saline (norm) imaging buffer, followed by acidic buffer (pH 5.5) which quenches cell surface fluorescence (i.e. sypHy localised to plasma membrane) to reveal background autofluorescence. Neurons were washed with saline imaging buffer before being perfused with NH_4_Cl buffer. Values are mean ± SEM. **F**: Scatter plots shows % vesicular sypHy ([(total – background) – (surface – background)] / (total – background)) x100) averaged over individual coverslips. n = 18 - 20 individual fields of view from independent coverslips, mean ± SEM indicated. Forest plot shows estimated difference between genotype with 95% CI, Welch’s t-test, p = 0.2674.

### Loss of Nlgn1 does not affect size of vesicle pools

Previous studies have demonstrated that the neurexin-Nlgn1 complex may regulate synaptic vesicle pools (Wittenmayer et al., 2009;Quinn et al., 2017;van Stegen et al., 2017). To examine whether loss of Nlgn1 impacts the size of distinct synaptic vesicle pools, WT and *Nlgn1*^-/-^ neurons expressing sypHy were subjected to two different trains of electrical field stimulation to mobilise the readily-releasable pool (60AP, 30Hz) and the reserve pool (1200AP, 10Hz), which together comprise the recycling pool of vesicles. When assayed in the presence of bafilomycin, the cumulative increase in sypHy fluorescence induced by each stimulation paradigm, as a proportion of total fluorescence revealed by NH4Cl, provides a measure of the relative amount of synaptic vesicles occupying each pool (Fig 1C). The remaining vesicles, which do not undergo activity-dependent fusion and thus are not part of the recycling pool, comprise the resting pool. The size of the readily releasable, reserve and resting pools are similar in both WT and *Nlgn1*^-/-^ neurons (Fig 1D, two-way ANOVA with Sidak’s multiple comparison test; readily releasable pool, WT = 24.16 ± 1.417 % total sypHy fluorescence, *Nlgn1*^-/-^ = 20.73 ± 1.296 %, p = 0.4256; reserve pool, WT = 36.06 ± 1.742, *Nlgn1*^-/-^ = 33.86 ± 2.056, p = 0.7539; resting pool, WT = 39.78 ± 1.639, *Nlgn1*^-/-^ = 45.41 ± 2.154, p = 0.0744). Thus, Nlgn1 does not act in a transsynaptic manner to modulate the partitioning of vesicles to different pools.

We also examined whether there was any difference in the total number of vesicles present in nerve terminals from *Nlgn1*^-/-^ and WT neurons. To investigate this, we looked at the partitioning of sypHy between different membranous components by perfusing neurons with acidic buffer (pH 5.5) to quench any surface pHluorin fluorescence thus leaving only background fluorescence, and then with NH4Cl to reveal total pHluorin fluorescence (Fig 1D). This allowed us to assess vesicular fluorescence as a proportion of total sypHy fluorescence, which was not changed for *Nlgn1*^-/-^ neurons compared to WT (Fig 1F; WT = 59.75 ± 3.561 % total sypHy fluorescence, *Nlgn1*^-/-^ = 64.20 ± 1.265, p = 0.2674 unpaired t-test). There was also no difference in total sypHy fluorescence in *Nlgn1*^-/-^ and WT neurons (Fig 1B; WT = 4686 ± 300.8 AU, *Nlgn1*^-/-^ = 4598 ± 394.4 AU, p = 0.8601 unpaired t-test). Together, this demonstrates that absence of Nlgn1 has no impact on the total number of vesicles in nerve terminals.

### Loss of Nlgn1 causes a change in the balance of stimulation-induced exocytosis and endocytosis

Having established that loss of Nlgn1 does not affect vesicle pools, we next sought to ascertain whether Nlgn1 has any impact on the dynamic cycling of vesicles. To examine this, we stimulated WT or *Nlgn1*^-/-^ neurons with 300 action potentials (10Hz) in the absence of bafilomycin and measured changes in sypHy fluorescence (Fig 2A). We observed that *Nlgn1*^-/-^ neurons displayed a dramatic reduction in the peak magnitude of pHluorin fluorescence measured during stimulation in this assay (Fig 2B, WT = 0.5383 ± 0.04042 maximal evoked fluorescence, *Nlgn1*^-/-^ = 0.3919 ± 0.02530, p = 0.0055 unpaired t-test). The peak height in this assay is a measure of net change in fluorescence during stimulation, and reflects a change in the equilibrium between exocytosis and endocytosis of synaptic vesicles during the stimulus train; thus a change in either of these parameters can lead to a change in peak height.

**Figure 2:**
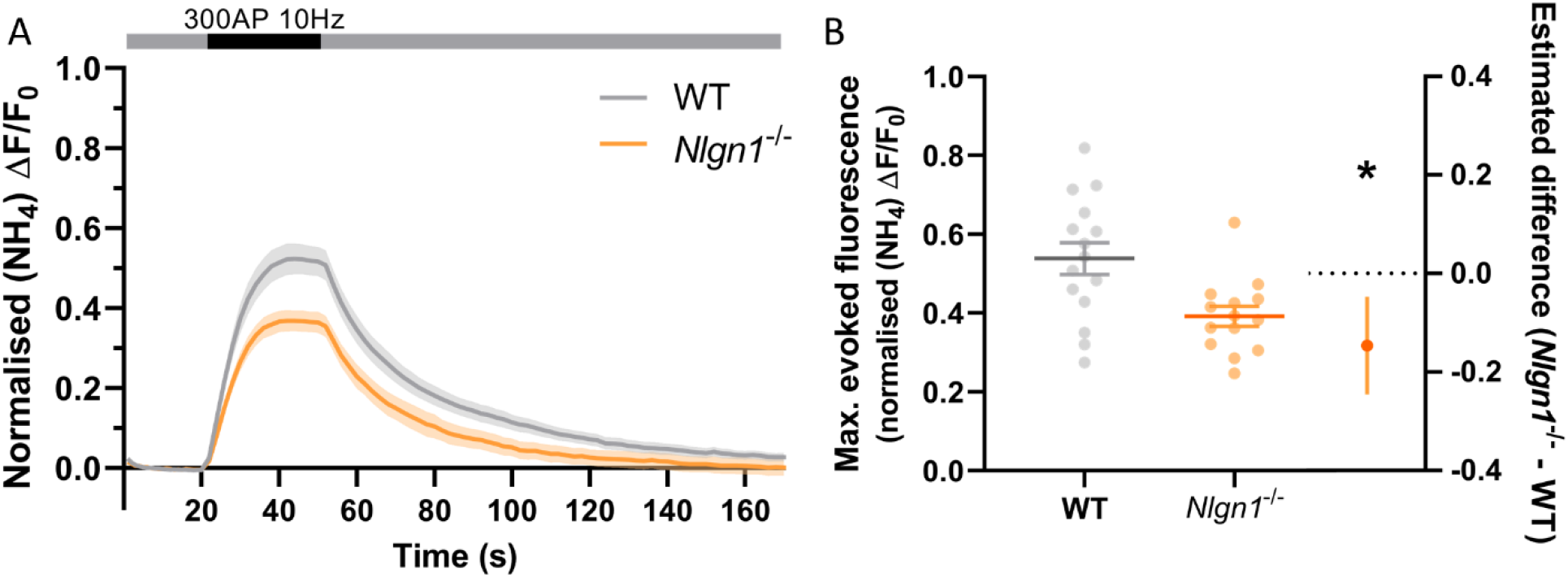
*Nlgn1*^-/-^ synapses display a change in the balance of exocytosis and endocytosis. *Nlgn1*^-/-^ and WT neurons transfected with sypHy were stimulated with 300 action potentials at 10Hz, and change in fluorescence monitored over time. Neurons were then perfused with NH4Cl to reveal total sypHy fluorescence. **A**: Time trace of mean (± SEM) changes in sypHy fluorescence, normalised to total fluorescence. **B**: Peak sypHy fluorescence during stimulus train, normalised to total fluorescence. Scatter plots show individual n, bars indicate mean ± SEM. Forest plot shows estimated difference between group means with 95% CI, unpaired t-test, \**p* < 0.05 (p = 0.0055). N = 14-15 individual fields of view from independent coverslips.

### Change in equilibrium between exocytosis and endocytosis in Nlgn1^-/-^ neurons is not caused by slower rate of exocytosis

To examine whether a slowing in the rate of exocytosis was responsible for the decrease in net change in evoked sypHy fluorescence in *Nlgn1*^-/-^ neurons (Fig2), we again stimulated the neurons (1200AP, 10Hz) in the presence of bafilomycin to prevent vesicle reacidification (Fig 3A) and monitored the change in pHluorin fluorescence, which purely reflects exocytosis of vesicles. Intriguingly, there was no global difference in exocytosis between *Nlgn1*^-/-^ and WT neurons (Fig 3A). Similarly, there was no change in the rate of exocytosis, either over the entire length of the stimulus train (measured as tau, Fig 3B, τ WT = 38.47 ± 5.86 s, *Nlgn1*^-/-^ = 33.91 ± 6.16 s, p = 0.5997, unpaired t-test) or over the initial portion of the stimulus train where the rate is largely linear (Fig 3C; Fig 3D WT = 0.02339 ± 0.002434 s^-1^, *Nlgn1*^-/-^ = 0.02173 ± 0.002124 s^-1^, p = 0.6186, unpaired t-test). We then normalised the traces to total sypHy fluorescence revealed by perfusion with NH4Cl, to ascertain the proportion of vesicles mobilised by stimulus train (Fig 3E). We found no change to the total recycling pool of vesicles (Fig 3F, WT = 59.46 ± 3.658 % total vesicles, *Nlgn1*^-/-^ = 59.84 ± 3.924 %, p = 0.9445 by unpaired t-test), in line with our earlier observation of lack of Nlgn1 having no effect on the readily releasable and reserve pools (Fig 1D). These data indicate that loss of Nlgn1 does not change the efficiency of exocytosis, and instead causes an increase in the rate of endocytosis during stimulation. Additionally, there was no change to the size of the recycling pool of vesicles.

**Figure 3:**
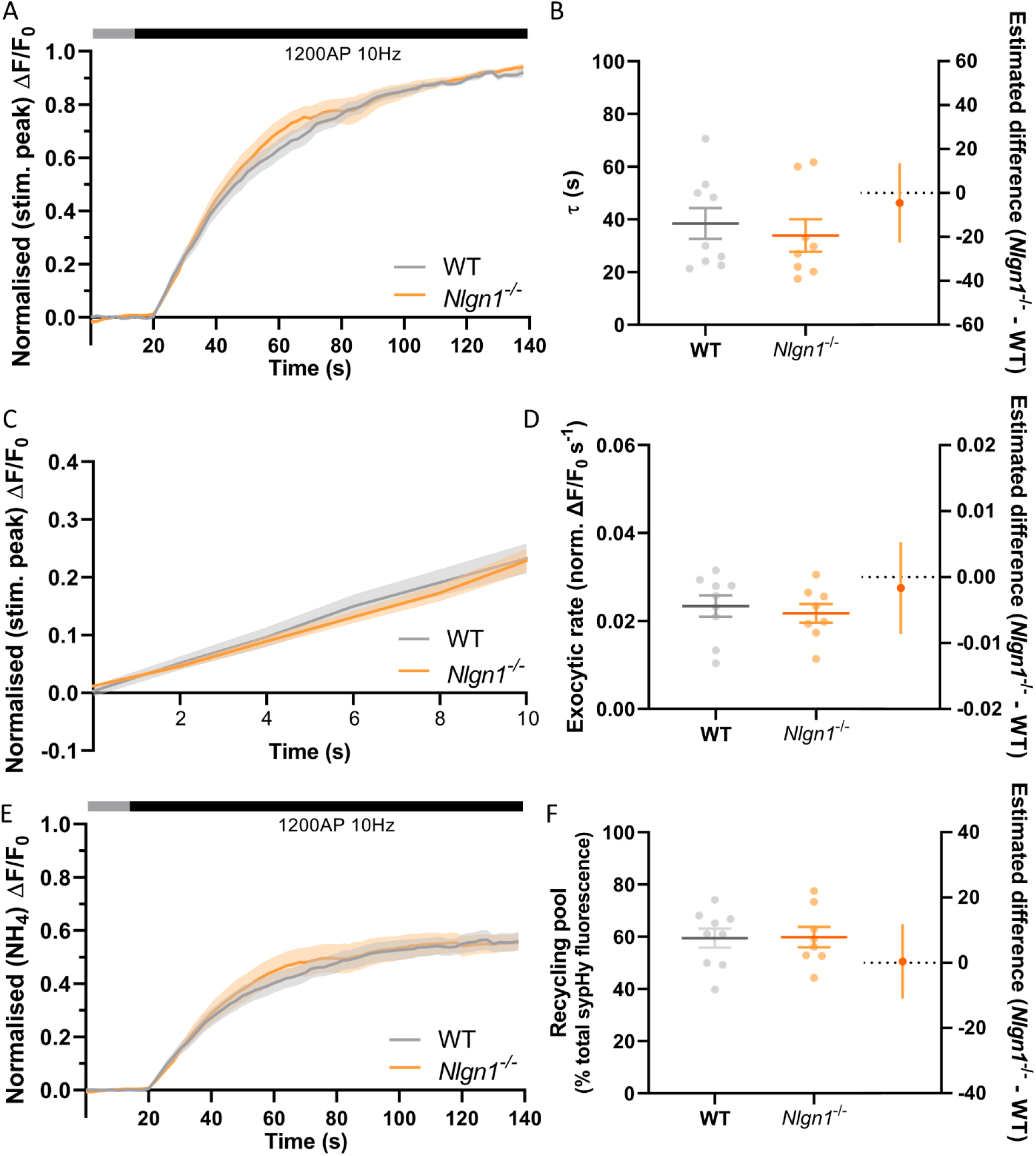
Rate of exocytosis is unchanged in *Nlgn1*^-/-^ synapses. *Nlgn1*^-/-^ and WT neurons transfected with sypHy were stimulated with 1200 AP (10Hz) in the presence of bafilomycin. **A**: Time trace of mean (± SEM) change in sypHy fluorescence, normalised to peak fluorescence during stimulation. **B**: Mean (± SEM) exocytosis time constant (tau, s), p = 0.5997. **C**: Time trace of mean (± SEM) change in sypHy fluorescence during first 10s of stimulation, normalised to peak fluorescence during stimulation. **D**: Mean (± SEM) rate of exocytosis (ΔF/F0 s^-1^) as assessed by linear fit over first 10 s of stimulation (from C), p = 0.6186. **E**: Time trace of mean (± SEM) change in sypHy fluorescence, normalised to total fluorescence revealed by NH4Cl perfusion. **F**: Mean (± SEM) size of total recycling pool assessed as peak fluorescence during stimulus train, as a % of total sypHy fluorescence revealed by NH4Cl, p = 0.9445. **B, D, F**: Scatter plots show individual data points. Bars indicate mean ± SEM. Forest plots show estimated difference between genotypes with 95% CI, unpaired t-test. N = 8-9 individual fields of view from independent coverslips.

### Loss of Nlgn1 specifically impacts compensatory endocytosis during stimulation

*Nlgn1*^-/-^ neurons have an increased rate of endocytosis during the stimulus train which leads to a decrease in the peak height of sypHy fluorescence during synchronous evoked release. To determine whether these changes to the rate of endocytosis persist following cessation of action potential stimulation, we examined the decay in pHluorin fluorescence following 300AP (10Hz) stimulation. Traces normalised to the peak fluorescence during stimulation revealed there was no difference in sypHy fluorescence between WT and *Nlgn1*^-/-^ synapses across any time points (Fig 4A, same data as Fig.2A normalised to peak of stimulation, mixed model ANOVA with Sidak’s multiple comparison test, p > 0.05 across all time points). We then performed exponential curve fitting to the post-stimulus portion of the sypHy trace, and confirmed that there was no change to the rate of compensatory endocytosis following the cessation of the stimulus train). Fig 4B, τ WT = 27.31 ± 1.70, *Nlgn1*^-/-^ = 23.78 ± 2.38, p = 0.2332, unpaired t-test). Thus, absence of Nlgn1 results in an increase in the rate of activity-dependent endocytosis specifically during neuronal activity.

**Figure 4:**
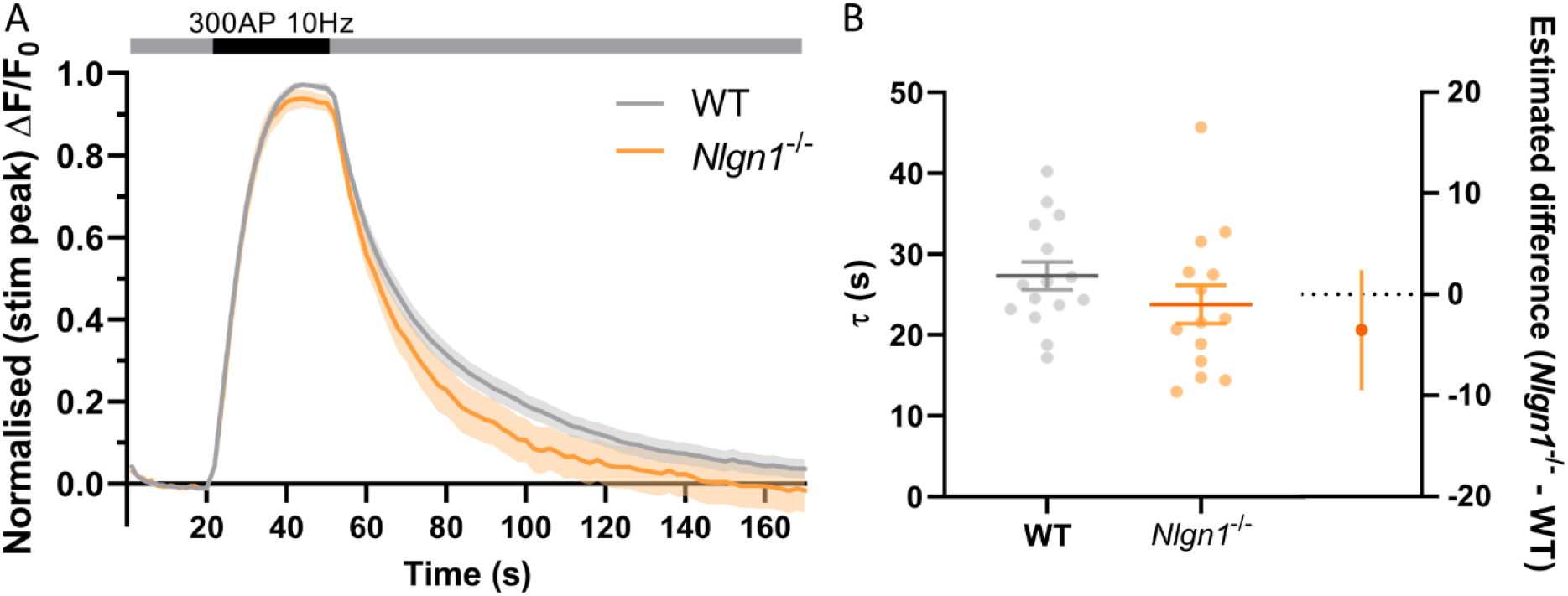
Loss of *Nlgn1*^-/-^ does not impact post-stimulus compensatory endocytosis. *Nlgn1*^-/-^ and WT neurons transfected with sypHy were stimulated with 300 action potentials at 10Hz, and change in fluorescence monitored over time. **A**: Time trace of mean (± SEM) changes in sypHy fluorescence, normalised to peak fluorescence during stimulus train. Mixed model ANOVA with Sidak’s multiple comparison test, p > 0.05 across all time points. **B**: Mean (± SEM) endocytic decay time constant (tau, s) from single exponential decay curve fit to data following end of stimulus train (from fig 2A). Scatter plots show individual n (single coverslip average), bars indicate mean ± SEM. Forest plot shows estimated difference between group means with 95% CI, unpaired t-test, *p* = 0.2332. N = 14-15 individual fields of view from independent coverslips, as in Figure 2.

## Discussion

In this study, through specific examination of presynaptic activity independent of postsynaptic signalling, we have revealed that the loss of Nlgn1 has transsynaptic impacts on synaptic vesicle cycling. Contrary to expectations, loss of Nlgn1 has no effect on the size of the readily releasable or recycling pool of synaptic vesicles. Instead, there was a large decrease in the change in pHluorin fluorescence during electrical field stimulation in *Nlgn1*^-/-^ neurons, leading to a lower stimulation-induced peak height. This was not driven by an altered rate of exocytosis, and thus likely reflects an increase in the rate of activity-dependent endocytosis and pHluorin retrieval to synaptic vesicles during neuronal activity, which is not accompanied by a change in the rate of compensatory endocytosis following the cessation of synchronous evoked neurotransmitter release. Thus, this is the first study to reveal that a cell adhesion protein that is localised exclusively to the postsynaptic compartment alters presynaptic endocytosis.

Synaptic vesicle endocytosis occurs on multiple time scales and encompasses several different pathways which have different retrieval capacities, are controlled by different molecular mediators, and may be spatiotemporally separable (Kononenko and Haucke, 2015;Cousin, 2017;Watanabe and Boucrot, 2017;Chanaday and Kavalali, 2018). Ultrafast endocytosis occurs over a 50-500ms time frame at physiological temperatures (Watanabe et al., 2013) and contributes a relatively small fraction of vesicle retrieval during trains of activity. The majority of membrane retrieval is driven by alternate forms of endocytosis (Soykan et al., 2017), including activity-dependent bulk endocytosis, which is strongly linked to neuronal activity, such that its activity dissipates immediately after the cessation of a stimulus train (Clayton and Cousin, 2009). Other forms of endocytosis, including clathrin-mediated and clathrin-independent compensatory endocytosis, are active both during and after the end of neuronal depolarisation. The transsynaptic complex formed by Nlgn1 may be important for shaping the endocytic pathways during neuronal activity, such that loss of Nlgn1 leads to a faster mode of vesicle recycling (such as ultrafast or bulk endocytosis) becoming dominant.

Intriguingly, other transsynaptic and cell adhesion proteins have also been directly shown to influence presynaptic endocytosis. Brain-specific deletion of the immunoglobulin superfamily cell adhesion protein CAR (coxsackievirus and adenovirus receptor; CAR-CNS^KO^ mice), which, similarly to Nlgn1 interacts with a range of postsynaptic PDZ-containing proteins including PSD95, also results in an increase in the rate of endocytosis (Wrackmeyer et al., 2019). However, unlike Nlgn1, CAR however is found in both presynaptic and postsynaptic compartments (Wrackmeyer et al., 2019). N-cadherin is a transsynaptic scaffold protein extracellularly localised between the pre-and post-synaptic membranes, at the edge of the active zone (Uchida et al., 1996), though it has also been observed more centrally at the extracellular side of the active zone, where it becomes enriched upon synaptic activity (Yam et al., 2013). Loss of N-cadherin specifically impaired post-stimulus compensatory endocytosis in nerve terminals which were subjected to intense stimulus trains (800AP at 20Hz, but not 200AP at 20Hz) which exhibited a high degree of exocytosis (van Stegen et al., 2017), and recent studies suggest that both synaptic vesicle and bulk endocytosis is impaired when total or postsynaptic N-cadherin is depleted (under physiological temperatures) (preprint; DOI: 10.1101/2021.02.05.429924). Conversely, overexpression of cadherin increased the rate of post-stimulus compensatory endocytosis (van Stegen et al., 2017).

Our study thus adds to growing evidence that transsynaptic scaffolding and cell adhesion proteins are important for controlling activity-induced endocytosis in presynaptic neurons, amongst their other synaptic activities. N-cadherin and/or CAR may modulate post-stimulus endocytosis, whilst neurexin and neuroligin may help define the prevalent modes of endocytosis during stimulation. Indeed, the post-stimulus rate of compensatory endocytosis has previously been shown to be unaffected by overexpression of Nlgn1 (van Stegen et al., 2017). Previous work has consistently demonstrated that Nlgn1 is required for inducing long-term potentiation (LTP) in the postsynaptic terminal via high frequency (100Hz) stimulus trains (Blundell et al., 2010;Jiang et al., 2017;Wu et al., 2019). Our findings raise the possibility that control of presynaptic recycling of synaptic vesicles may contribute to Nlgn1-dependent LTP. Synaptic vesicles that are generated by different recycling modes undergo different sorting pathways across different timescales, and repopulate distinct vesicle pools with different release probabilities and synaptic localisation (Kokotos and Cousin, 2015;Rey et al., 2020;Ivanova et al., 2021). Thus, changes in endocytosis may alter the firing pattern of individual and groups of neurons during sustained activity *in vivo* thereby altering the information they encode.

How might transsynaptic protein complexes and signalling contribute to the coding of presynaptic vesicle recycling modalities? A C-terminally deleted version of Nlgn1, which is unable to bind PDZ domain containing PSD proteins, altered the alignment of the presynaptic active zone with post-synaptic receptors in a dominant-negative manner, impairing neurotransmission (Haas et al., 2018). Thus, we know that Nlgn1 can act in a transsynaptic manner to alter the presynaptic molecular architecture, likely via interactions with neurexin. Neurexin controls clustering and function of presynaptic Ca^2+^ channels (Gandini and Zamponi, 2021), perhaps via changing the mobility of α2δ-1 auxiliary subunits (Brockhaus et al., 2018). There is strong evidence that Ca^2+^ influx can mediate the speed of multiple forms of endocytosis, including fast forms of endocytosis (Wu et al., 2005;Kononenko and Haucke, 2015;Morton et al., 2015), though exactly how Ca^2+^ affects distinct modes of endocytosis in specific synapses remains unclear (Leitz and Kavalali, 2016). Neurexin also regulates F-actin assembly (Rui et al., 2017), which is essential for multiple vesicle retrieval pathways, including ultrafast and bulk endocytosis (Watanabe et al., 2014;Wu et al., 2016;Soykan et al., 2017). Thus, Nlgn1 may act via neurexin to control local presynaptic F-actin stability or localise Ca^2+^ microdomains, thus impacting endocytic pathways. Future work is required to ascertain the molecular mechanisms by which Nlgn1 impacts endocytosis, and indeed, which endocytic pathways are altered by the absence of Nlgn1.

At the systems level, an interesting question is whether disruptions to synaptic molecules involved in endocytosis lead to the manifestation of overlapping behavioural symptoms. Human mutations in the neuroligin gene family have been repeatedly documented in neurodevelopmental disorders, including several *NLGN1* variants identified in autism spectrum disorders, of which the majority are loss-of-function mutations reducing Nlgn1 expression (ASD)(Vieira et al., 2021). Indeed, gene mutations in various proteins that regulate endocytosis have been identified in neurodevelopmental disorders, predominantly epilepsy and intellectual disability (Bonnycastle et al., 2020;John et al., 2021), including proteins such as VAMP4 which has specific roles in activity-dependent bulk endocytosis (Monies et al., 2017). This further strengthens the growing evidence that dysfunctions at the synapse represents a core convergent pathway that underlies disease mechanisms in these neurodevelopmental disorders or synaptopathies (Grant, 2012).

However, behavioural and cognitive analyses of mouse models reveal a complexity in behavioural disturbances. Previous work indicated mice lacking Nlgn1 showed no changes across numerous behavioural measures including anxiety-like behaviours in the elevated plus maze, but displayed impaired spatial learning and memory in the Morris Water Maze (Blundell et al., 2010). We have recently shown, through a deep cognitive characterization using a series of touchscreen-based assays, that while associative learning was not impacted due to loss of Nlgn1, motivated behaviour and cost-reward processing was consistently disrupted (Luo et al., 2020), having implications for mood disorders. Whilst postsynaptic mechanisms are likely a major driver to this altered behaviour, changes to endocytosis may also contribute to these effects. Unsurprisingly, mouse models involving disruption to core presynaptic endocytosis machinery such as dynamin, synaptojanin, or auxilin often lead to early lethality or seizures (Cremona et al., 1999;Ferguson et al., 2007;Boumil et al., 2010;Yim et al., 2010;Luo et al., 2017). However, other models, where endocytosis is partially compromised but not ablated, exhibit varied alterations in exploratory and locomotor behaviour, as well as learning and memory. For example, amphiphysin 1^-/-^, endophilin A1^-/-^, or intersectin1 mutant mice displayed learning deficits in the Morris water maze and reduced contextual fear memory (Di Paolo et al., 2002;Sengar et al., 2013;Yang et al., 2018). More detailed investigations into the contribution of different molecular players at both the pre- and postsynapse to the regulation of distinct components of behaviour and cognition will help reveal how integration of these factors controls complex behaviour in health and disease.

Synaptic adhesion molecules, including the neurexin-neuroligin complex, have been theorised to define the organisation and specialisation, or coding, of synapses (Sudhof, 2021). Perhaps a key distinguishing property of this synaptic code is the capacity to sustain specific trains of stimulation frequencies. Thus, control of the modality by which synapses reset and recycle vesicles may be an important function of these proteins, which would have knock-on effects on neural circuit activity and complex behaviours.

## Conflict of interest

The authors declare no conflict of interest.

## Data availability

Datasets are available on request; the raw data supporting the conclusions of this article will be made available by the authors, without undue reservation.

## Author Contributions

J.K.L., J.N. and S.L.G. contributed to conception and design of the study. J.K.L. performed experiments with training support from H.M. J.K.L. and H.M. analysed the data. J.K.L. performed statistical analysis. S.L.G. and J.K.L. wrote the first draft of the manuscript. All authors contributed to manuscript revision, read, and approved the submitted version. J.N. and S.L.G. acquired funding support for the project.

## Funding

Funding: This work was supported by an Australian Research Council Future Fellowship (FT140101327) to J.N., a National Health and Medical Research Council of Australia Career Development Fellowship (GNT1105478) and Florey Fellowship to S.L.G., an Australian Government Research Training Program Scholarship to J.L and H.M, and National Health and Medical Research Council of Australia Project Grants to J.N. (GNT1083334) and S.L.G., (GNT1085483). The Florey Institute of Neuroscience and Mental Health acknowledges the strong support from the Victorian Government and in particular the funding from the Operational Infrastructure Support Grant.

## Acknowledgements

The authors thank the Core Animal Services staff and Bioresources Facilities of the Florey Institute for assistance with animal maintenance, the Florey Neuroscience Microscopy Facility for their support and expert technical assistance, and Linda Nguyen of Cell to Sketch for creating Figure 1A. The authors thank the Gordon and Nithianantharajah labs for useful discussions.

## Notes

### Competing Interest Statement

The authors have declared no competing interest.

## References

Arac, D., Boucard, A.A., Ozkan, E., Strop, P., Newell, E., Sudhof, T.C., and Brunger, A.T. (2007). Structures of neuroligin-1 and the neuroligin-1/neurexin-1 beta complex reveal specific protein-protein and protein-Ca2+ interactions. Neuron 56, 992–1003.

Blundell, J., Blaiss, C.A., Etherton, M.R., Espinosa, F., Tabuchi, K., Walz, C., Bolliger, M.F., Sudhof, T.C., and Powell, C.M. (2010). Neuroligin-1 deletion results in impaired spatial memory and increased repetitive behavior. J Neurosci 30, 2115–2129.

Bonnycastle, K., Davenport, E.C., and Cousin, M.A. (2020). Presynaptic dysfunction in neurodevelopmental disorders: Insights from the synaptic vesicle life cycle. J Neurochem.

Boumil, R.M., Letts, V.A., Roberts, M.C., Lenz, C., Mahaffey, C.L., Zhang, Z.W., Moser, T., and Frankel, W.N. (2010). A missense mutation in a highly conserved alternate exon of dynamin-1 causes epilepsy in fitful mice. PLoS Genet 6.

Brockhaus, J., Schreitmuller, M., Repetto, D., Klatt, O., Reissner, C., Elmslie, K., Heine, M., and Missler, M. (2018). alpha-Neurexins Together with alpha2delta-1 Auxiliary Subunits Regulate Ca(2+) Influx through Cav2.1 Channels. J Neurosci 38, 8277–8294.

Budreck, E.C., Kwon, O.B., Jung, J.H., Baudouin, S., Thommen, A., Kim, H.S., Fukazawa, Y., Harada, H., Tabuchi, K., Shigemoto, R., Scheiffele, P., and Kim, J.H. (2013). Neuroligin-1 controls synaptic abundance of NMDA-type glutamate receptors through extracellular coupling. Proc Natl Acad Sci U S A 110, 725–730.

Chanaday, N.L., and Kavalali, E.T. (2018). Optical detection of three modes of endocytosis at hippocampal synapses. Elife 7.

Chubykin, A.A., Atasoy, D., Etherton, M.R., Brose, N., Kavalali, E.T., Gibson, J.R., and Sudhof, T.C. (2007). Activity-dependent validation of excitatory versus inhibitory synapses by neuroligin-1 versus neuroligin-2. Neuron 54, 919–931.

Clayton, E.L., and Cousin, M.A. (2009). The molecular physiology of activity-dependent bulk endocytosis of synaptic vesicles. J Neurochem 111, 901–914.

Cousin, M.A. (2017). Integration of Synaptic Vesicle Cargo Retrieval with Endocytosis at Central Nerve Terminals. Front Cell Neurosci 11, 234.

Cremona, O., Di Paolo, G., Wenk, M.R., Luthi, A., Kim, W.T., Takei, K., Daniell, L., Nemoto, Y., Shears, S.B., Flavell, R.A., Mccormick, D.A., and De Camilli, P. (1999). Essential role of phosphoinositide metabolism in synaptic vesicle recycling. Cell 99, 179–188.

Di Paolo, G., Sankaranarayanan, S., Wenk, M.R., Daniell, L., Perucco, E., Caldarone, B.J., Flavell, R., Picciotto, M.R., Ryan, T.A., Cremona, O., and De Camilli, P. (2002). Decreased synaptic vesicle recycling efficiency and cognitive deficits in amphiphysin 1 knockout mice. Neuron 33, 789–804.

Espinosa, F., Xuan, Z., Liu, S., and Powell, C.M. (2015). Neuroligin 1 modulates striatal glutamatergic neurotransmission in a pathway and NMDAR subunit-specific manner. Front Synaptic Neurosci 7, 11.

Ferguson, S.M., Brasnjo, G., Hayashi, M., Wolfel, M., Collesi, C., Giovedi, S., Raimondi, A., Gong, L.W., Ariel, P., Paradise, S., O’toole, E., Flavell, R., Cremona, O., Miesenbock, G., Ryan, T.A., and De Camilli, P. (2007). A selective activity-dependent requirement for dynamin 1 in synaptic vesicle endocytosis. Science 316, 570–574.

Gandini, M.A., and Zamponi, G.W. (2021). Voltage-gated calcium channel nanodomains: molecular composition and function. FEBS J.

Gordon, S.L., Leube, R.E., and Cousin, M.A. (2011). Synaptophysin is required for synaptobrevin retrieval during synaptic vesicle endocytosis. J Neurosci 31, 14032–14036.

Graf, E.R., Zhang, X., Jin, S.X., Linhoff, M.W., and Craig, A.M. (2004). Neurexins induce differentiation of GABA and glutamate postsynaptic specializations via neuroligins. Cell 119, 1013–1026.

Grant, S.G. (2012). Synaptopathies: diseases of the synaptome. Curr Opin Neurobiol 22, 522–529.

Haas, K.T., Compans, B., Letellier, M., Bartol, T.M., Grillo-Bosch, D., Sejnowski, T.J., Sainlos, M., Choquet, D., Thoumine, O., and Hosy, E. (2018). Pre-post synaptic alignment through neuroligin-1 tunes synaptic transmission efficiency. Elife 7.

Heine, M., Thoumine, O., Mondin, M., Tessier, B., Giannone, G., and Choquet, D. (2008). Activity-independent and subunit-specific recruitment of functional AMPA receptors at neurexin/neuroligin contacts. Proc Natl Acad Sci U S A 105, 20947–20952.

Ichtchenko, K., Hata, Y., Nguyen, T., Ullrich, B., Missler, M., Moomaw, C., and Sudhof, T.C. (1995). Neuroligin 1: a splice site-specific ligand for beta-neurexins. Cell 81, 435–443.

Irie, M., Hata, Y., Takeuchi, M., Ichtchenko, K., Toyoda, A., Hirao, K., Takai, Y., Rosahl, T.W., and Sudhof, T.C. (1997). Binding of neuroligins to PSD-95. Science 277, 1511–1515.

Ivanova, D., Dobson, K.L., Gajbhiye, A., Davenport, E.C., Hacker, D., Ultanir, S.K., Trost, M., and Cousin, M.A. (2021). Control of synaptic vesicle release probability via VAMP4 targeting to endolysosomes. Sci Adv 7.

Jiang, M., Polepalli, J., Chen, L.Y., Zhang, B., Sudhof, T.C., and Malenka, R.C. (2017). Conditional ablation of neuroligin-1 in CA1 pyramidal neurons blocks LTP by a cell-autonomous NMDA receptor-independent mechanism. Mol Psychiatry 22, 375–383.

John, A., Ng-Cordell, E., Hanna, N., Brkic, D., and Baker, K. (2021). The neurodevelopmental spectrum of synaptic vesicle cycling disorders. J Neurochem 157, 208–228.

Jung, S.Y., Kim, J., Kwon, O.B., Jung, J.H., An, K., Jeong, A.Y., Lee, C.J., Choi, Y.B., Bailey, C.H., Kandel, E.R., and Kim, J.H. (2010). Input-specific synaptic plasticity in the amygdala is regulated by neuroligin-1 via postsynaptic NMDA receptors. Proc Natl Acad Sci U S A 107, 4710–4715.

Kim, J., Jung, S.Y., Lee, Y.K., Park, S., Choi, J.S., Lee, C.J., Kim, H.S., Choi, Y.B., Scheiffele, P., Bailey, C.H., Kandel, E.R., and Kim, J.H. (2008). Neuroligin-1 is required for normal expression of LTP and associative fear memory in the amygdala of adult animals. Proc Natl Acad Sci U S A 105, 9087–9092.

Kokotos, A.C., and Cousin, M.A. (2015). Synaptic vesicle generation from central nerve terminal endosomes. Traffic 16, 229–240.

Kononenko, N.L., and Haucke, V. (2015). Molecular mechanisms of presynaptic membrane retrieval and synaptic vesicle reformation. Neuron 85, 484–496.

Leitz, J., and Kavalali, E.T. (2016). Ca2+ Dependence of Synaptic Vesicle Endocytosis. Neuroscientist 22, 464–476.

Luo, J., Norris, R.H., Gordon, S.L., and Nithianantharajah, J. (2017). Neurodevelopmental synaptopathies: Insights from behaviour in rodent models of synapse gene mutations. Prog Neuropsychopharmacol Biol Psychiatry.

Luo, J., Tan, J.M., and Nithianantharajah, J. (2020). A molecular insight into the dissociable regulation of associative learning and motivation by the synaptic protein neuroligin-1. BMC Biol 18, 118.

Missler, M., Zhang, W., Rohlmann, A., Kattenstroth, G., Hammer, R.E., Gottmann, K., and Sudhof, T.C. (2003). Alpha-neurexins couple Ca2+ channels to synaptic vesicle exocytosis. Nature 423, 939–948.

Mondin, M., Labrousse, V., Hosy, E., Heine, M., Tessier, B., Levet, F., Poujol, C., Blanchet, C., Choquet, D., and Thoumine, O. (2011). Neurexin-neuroligin adhesions capture surface-diffusing AMPA receptors through PSD-95 scaffolds. J Neurosci 31, 13500–13515.

Monies, D., Abouelhoda, M., Alsayed, M., Alhassnan, Z., Alotaibi, M., Kayyali, H., Al-Owain, M., Shah, A., Rahbeeni, Z., Al-Muhaizea, M.A., Alzaidan, H.I., Cupler, E., Bohlega, S., Faqeih, E., Faden, M., Alyounes, B., Jaroudi, D., Goljan, E., Elbardisy, H., Akilan, A., Albar, R., Aldhalaan, H., Gulab, S., Chedrawi, A., Al Saud, B.K., Kurdi, W., Makhseed, N., Alqasim, T., El Khashab, H.Y., Al-Mousa, H., Alhashem, A., Kanaan, I., Algoufi, T., Alsaleem, K., Basha, T.A., Al-Murshedi, F., Khan, S., Al-Kindy, A., Alnemer, M., Al-Hajjar, S., Alyamani, S., Aldhekri, H., Al-Mehaidib, A., Arnaout, R., Dabbagh, O., Shagrani, M., Broering, D., Tulbah, M., Alqassmi, A., Almugbel, M., Alquaiz, M., Alsaman, A., Al-Thihli, K., Sulaiman, R.A., Al-Dekhail, W., Alsaegh, A., Bashiri, F.A., Qari, A., Alhomadi, S., Alkuraya, H., Alsebayel, M., Hamad, M.H., Szonyi, L., Abaalkhail, F., Al-Mayouf, S.M., Almojalli, H., Alqadi, K.S., Elsiesy, H., Shuaib, T.M., Seidahmed, M.Z., Abosoudah, I., Akleh, H., Alghonaium, A., Alkharfy, T.M., Al Mutairi, F., Eyaid, W., Alshanbary, A., Sheikh, F.R., Alsohaibani, F.I., Alsonbul, A., Al Tala, S., Balkhy, S., Bassiouni, R., Alenizi, A.S., Hussein, M.H., Hassan, S., Khalil, M., Tabarki, B., Alshahwan, S., Oshi, A., Sabr, Y., Alsaadoun, S., Salih, M.A., Mohamed, S., Sultana, H., Tamim, A., El-Haj, M., Alshahrani, S., Bubshait, D.K., Alfadhel, M., et al. (2017). The landscape of genetic diseases in Saudi Arabia based on the first 1000 diagnostic panels and exomes. Hum Genet 136, 921–939.

Morton, A., Marland, J.R., and Cousin, M.A. (2015). Synaptic vesicle exocytosis and increased cytosolic calcium are both necessary but not sufficient for activity-dependent bulk endocytosis. J Neurochem 134, 405–415.

Nguyen, T., and Sudhof, T.C. (1997). Binding properties of neuroligin 1 and neurexin 1beta reveal function as heterophilic cell adhesion molecules. J Biol Chem 272, 26032–26039.

Quinn, D.P., Kolar, A., Wigerius, M., Gomm-Kolisko, R.N., Atwi, H., Fawcett, J.P., and Krueger, S.R. (2017). Pan-neurexin perturbation results in compromised synapse stability and a reduction in readily releasable synaptic vesicle pool size. Sci Rep 7, 42920.

Rey, S., Marra, V., Smith, C., and Staras, K. (2020). Nanoscale Remodeling of Functional Synaptic Vesicle Pools in Hebbian Plasticity. Cell Rep 30, 2006–2017 e2003.

Rui, M., Qian, J., Liu, L., Cai, Y., Lv, H., Han, J., Jia, Z., and Xie, W. (2017). The neuronal protein Neurexin directly interacts with the Scribble-Pix complex to stimulate F-actin assembly for synaptic vesicle clustering. J Biol Chem 292, 14334–14348.

Sengar, A.S., Ellegood, J., Yiu, A.P., Wang, H., Wang, W., Juneja, S.C., Lerch, J.P., Josselyn, S.A., Henkelman, R.M., Salter, M.W., and Egan, S.E. (2013). Vertebrate intersectin1 is repurposed to facilitate cortical midline connectivity and higher order cognition. J Neurosci 33, 4055–4065.

Song, J.Y., Ichtchenko, K., Sudhof, T.C., and Brose, N. (1999). Neuroligin 1 is a postsynaptic cell-adhesion molecule of excitatory synapses. Proc Natl Acad Sci U S A 96, 1100–1105.

Soykan, T., Kaempf, N., Sakaba, T., Vollweiter, D., Goerdeler, F., Puchkov, D., Kononenko, N.L., and Haucke, V. (2017). Synaptic Vesicle Endocytosis Occurs on Multiple Timescales and Is Mediated by Formin-Dependent Actin Assembly. Neuron 93, 854–866 e854.

Sudhof, T.C. (2008). Neuroligins and neurexins link synaptic function to cognitive disease. Nature 455, 903–911.

Sudhof, T.C. (2021). The cell biology of synapse formation. J Cell Biol 220.

Uchida, N., Honjo, Y., Johnson, K.R., Wheelock, M.J., and Takeichi, M. (1996). The catenin/cadherin adhesion system is localized in synaptic junctions bordering transmitter release zones. J Cell Biol 135, 767–779.

Van Stegen, B., Dagar, S., and Gottmann, K. (2017). Release activity-dependent control of vesicle endocytosis by the synaptic adhesion molecule N-cadherin. Sci Rep 7, 40865.

Varoqueaux, F., Aramuni, G., Rawson, R.L., Mohrmann, R., Missler, M., Gottmann, K., Zhang, W., Sudhof, T.C., and Brose, N. (2006). Neuroligins determine synapse maturation and function. Neuron 51, 741–754.

Vieira, M.M., Jeong, J., and Roche, K.W. (2021). The role of NMDA receptor and neuroligin rare variants in synaptic dysfunction underlying neurodevelopmental disorders. Curr Opin Neurobiol 69, 93–104.

Watanabe, S., and Boucrot, E. (2017). Fast and ultrafast endocytosis. Curr Opin Cell Biol 47, 64–71.

Watanabe, S., Rost, B.R., Camacho-Perez, M., Davis, M.W., Sohl-Kielczynski, B., Rosenmund, C., and Jorgensen, E.M. (2013). Ultrafast endocytosis at mouse hippocampal synapses. Nature 504, 242–247.

Watanabe, S., Trimbuch, T., Camacho-Perez, M., Rost, B.R., Brokowski, B., Sohl-Kielczynski, B., Felies, A., Davis, M.W., Rosenmund, C., and Jorgensen, E.M. (2014). Clathrin regenerates synaptic vesicles from endosomes. Nature 515, 228–233.

Wittenmayer, N., Korber, C., Liu, H., Kremer, T., Varoqueaux, F., Chapman, E.R., Brose, N., Kuner, T., and Dresbach, T. (2009). Postsynaptic Neuroligin1 regulates presynaptic maturation. Proc Natl Acad Sci U S A 106, 13564–13569.

Wrackmeyer, U., Kaldrack, J., Juttner, R., Pannasch, U., Gimber, N., Freiberg, F., Purfurst, B., Kainmueller, D., Schmitz, D., Haucke, V., Rathjen, F.G., and Gotthardt, M. (2019). The cell adhesion protein CAR is a negative regulator of synaptic transmission. Sci Rep 9, 6768.

Wu, W., Xu, J., Wu, X.S., and Wu, L.G. (2005). Activity-dependent acceleration of endocytosis at a central synapse. J Neurosci 25, 11676–11683.

Wu, X., Morishita, W.K., Riley, A.M., Hale, W.D., Sudhof, T.C., and Malenka, R.C. (2019). Neuroligin-1 Signaling Controls LTP and NMDA Receptors by Distinct Molecular Pathways. Neuron 102, 621–635 e623.

Wu, X.S., Lee, S.H., Sheng, J., Zhang, Z., Zhao, W.D., Wang, D., Jin, Y., Charnay, P., Ervasti, J.M., and Wu, L.G. (2016). Actin Is Crucial for All Kinetically Distinguishable Forms of Endocytosis at Synapses. Neuron 92, 1020–1035.

Yam, P.T., Pincus, Z., Gupta, G.D., Bashkurov, M., Charron, F., Pelletier, L., and Colman, D.R. (2013). N-cadherin relocalizes from the periphery to the center of the synapse after transient synaptic stimulation in hippocampal neurons. PLoS One 8, e79679.

Yang, Y., Chen, J., Guo, Z., Deng, S., Du, X., Zhu, S., Ye, C., Shi, Y.S., and Liu, J.J. (2018). Endophilin A1 Promotes Actin Polymerization in Dendritic Spines Required for Synaptic Potentiation. Front Mol Neurosci 11, 177.

Yim, Y.I., Sun, T., Wu, L.G., Raimondi, A., De Camilli, P., Eisenberg, E., and Greene, L.E. (2010). Endocytosis and clathrin-uncoating defects at synapses of auxilin knockout mice. Proc Natl Acad Sci U S A 107, 4412–4417.

