## Supplementary Material for "Postsynaptic neuroligin-1 mediates presynaptic endocytosis during neuronal activity"

**Supplementary Methods**

**Quantifying size of synaptic vesicle pools**

To quantify the size of synaptic vesicle pools, the readily releasable and reserve pool were released sequentially by two separate trains of stimuli. Neurons were stimulated in the presence of 1 μM bafilomycin A1 with a train of 60 action potentials delivered at 30 Hz (100 mA, 1ms pulse width) to trigger fusion of the readily releasable pool, followed by 1200 action potentials at 10Hz to mobilise the reserve pool. Fluorescence change from baseline (ΔF) was calculated as (F – F0)/F0, where F0 is the baseline fluorescence prior to stimulation. ΔF is then normalized to the maximum (i.e. total) fluorescence change during alkaline imaging buffer perfusion (ΔF/F_total_). The maximum ΔF/F_total_ after the termination of the first stimulation and before the commencement of the second stimulation was used to report the size of the readily releasable pool. The maximum ΔF/F_total_ after the termination of the second stimulation and before alkaline imaging buffer perfusion was used to report the size of the total recycling pool. The size the reserve pool was calculated as total recycling pool – readily releasable pool.

**Monitoring dynamic cycling (exo- and endocytosis) of synaptic vesicles**

Neurons were stimulated with a train of 300 action potentials at 10 Hz (100 mA, 1ms pulse width), then imaged for a further 150 seconds to capture post-stimulus compensatory endocytosis (images captured at 2s intervals). Baseline subtracted fluorescence ΔF was normalised to maximum fluorescence revealed by NH_4_ to obtain ΔF/F_total_. The peak of ΔF/F_total_ during the stimulus train was taken as the stimulation peak. ΔF/F_total_ after the termination of stimulation and before perfusion with alkaline imaging buffer was fitted with the function:

$${\Delta F}/{F_{total}}=ae^{-kt}+plateau$$

 The parameter k represents endocytic rate.

**Quantifying exocytic rate and the size of the total recycling pool**

Neurons were stimulated with a train of 1200 action potentials at 10Hz (100 mA, 1ms pulse width) in the presence of 1 μM bafilomycin A1 to inhibit reacidification of synaptic vesicles. To estimate exocytic rate, ∆F was normalized to the peak fluorescence increase during stimulation (F_peak)_ to obtain ∆F /F_peak_ and fitted with the function:

${\Delta F}/{F_{peak}=1-e^{-kt}+F_{0}}$

The parameter k represents exocytic rate.

Early exocytic rate was estimated by fitting the ∆F/F_peak_ of the first five frames after stimulation commenced with the linear function:

$$\Delta F/F_{peak}=kt+F0$$

The parameter k represents early (linear) exocytic rate.

The maximum ∆F/F_total_ during the last 20 seconds (10 frames) prior to perfusion of alkaline imaging buffer expressed as a percentage was used quantify the size of the total recycling pool in this experiment.
